# Salicylate increases fitness cost associated with MarA-mediated antibiotic resistance

**DOI:** 10.1101/454983

**Authors:** T. Wang, C. Kunze, M. J. Dunlop

## Abstract

Antibiotic resistance is generally associated with a fitness deficit resulting from the burden of producing and maintaining resistance machinery. This additional cost suggests that resistant bacteria will be outcompeted by susceptible bacteria in conditions without antibiotics. However, in practice this process is slow due in part to regulation that minimizes expression of these genes in the absence of antibiotics. This suggests that if it were possible to turn on their expression, the cost would increase, thereby accelerating removal of resistant strains. Experimental and theoretical studies have shown that environmental chemicals can change the fitness cost associated with resistance and therefore have a significant impact on population dynamics. MarA (multiple antibiotic resistance activator) is a clinically important regulator in *Escherichia coli* which activates downstream genes to increase resistance against multiple classes of antibiotics. Salicylate is an inducer of MarA which can be found in the environment and de-represses *marA*’s expression. In this study, we sought to unravel the interplay between salicylate and the fitness cost of MarA-mediated antibiotic resistance. Using salicylate as a natural inducer of MarA, we found that a wide spectrum of concentrations can increase burden in resistant strains compared to susceptible strains. Induction resulted in rapid exclusion of resistant bacteria from mixed populations of antibiotic resistant and susceptible cells. A mathematical model captures the process and predicts its effect in various environmental conditions. Our work provides a quantitative understanding of salicylate exposure on the fitness of different MarA variants, and suggests that salicylate can lead to selection against MarA-mediated resistant strains. More generally, our findings show that natural inducers may serve to bias population membership and could impact antibiotic resistance and other important phenotypes.

## Introduction

Antibiotic resistance is frequently associated with fitness deficits such as those resulting from burdensome expression of resistance proteins or from excessive energy consumption by resistance machinery (1–5). These observations suggest that in the absence of antibiotic pressure, a bacterial population should be biased away from drug resistance (6–8). In practice, compensatory mutations and precise regulation of burdensome protein expression reduce the effective cost, largely eliminating the fitness differences. For example, in an antibiotic-free environment, tetracycline resistant bacteria that express the costly TetA efflux pump are expected to be outcompeted by susceptible strains (9). However, experiments have shown that the fitness cost of tetracycline resistance without antibiotic pressure is minimal (7, 10). Tight repression of *tetA* by TetR keeps costs low in the absence of inducers (7, 9). These regulatory approaches to reducing cost are common, and other examples include repression of the multiple antibiotic resistance activator *marA* by MarR (11) and the multi-drug efflux pump *acrAB* by AcrR (12). However, certain environmental chemicals can serve as inducers that relieve repression, and may change the fitness of antibiotic resistant bacteria.

Environmental inducers include both the substrates that these mechanisms protect against (e.g. antibiotics) and other compounds. For example, the tetracycline decay product anhydrotetracycline (aTc) (10), which is found widely in soil and wastewater (13, 14), significantly increases the fitness cost of tetracycline resistant cells by releasing repression by TetR. The presence of aTc will select against resistant cells in the absence of antibiotic pressure because TetA is costly, but provides no benefit (9, 10). Other environmental chemicals such as food preservatives (15) and pharmaceutical products (1) can impose fitness disadvantages by inducing higher levels of expression of resistance genes, which may lead to their eventual loss. A recent study demonstrated this effect by growing *E. coli* for 2000 generations in the presence of benzoate, an inducer for multiple antibiotic resistance genes. The majority of benzoate-evolved strains acquired mutations in *marA* and its homolog *rob*, thereby lowering resistance against chloramphenicol and tetracycline (15).

MarA is a clinically important antibiotic resistance regulator conserved across enteric bacteria (16). It increases resistance levels by regulating over 40 downstream genes (17–23), including multi-drug efflux pumps and porins (24, 25). Activation of these genes is taxing, resulting in decreased growth and a reduced fraction of cells with elevated MarA in a mixed community (1, 3). However, *marA* is under the control of its repressor MarR, which minimizes the cost in the absence of induction (21). MarA can also be modulated by its environmental inducer salicylate (1, 17), which binds directly with MarR and prevents repression of *marA* (16), significantly decreasing growth (1), which suggests an intriguing interplay between this environmental inducer and the cost of antibiotic machinery.

In this work, we focused on the cost of MarA-mediated resistance under salicylate exposure. We found that a wide range of concentrations of salicylate can induce MarA-mediated burden. AcrAB-TolC efflux pumps were found to be a major contributor to the salicylate-induced burden. The difference in cost under salicylate exposure leads to rapid exclusion of resistant strains in competitive settings. A mathematical model captures the process by which populations are biased away from resistance and predicts its effect in various salicylate conditions. This work suggests that the fitness cost of MarA-mediated antibiotic resistance can be amplified by salicylate, biasing populations towards susceptible strains.

## Materials and Methods

### Strains

We used four strains in this research: WT, MarA^−^, MarA^+^, and AcrB^−^. In the MarA^−^ strain, we deleted the *marRAB* operon, *rob* gene, and *soxSR* genes in *E. coli* MG1655 (26). In the MarA^+^ strain, we performed transversion mutations on MarR binding sites (23) in the chromosomal *marRAB* promoter to inhibit binding of MarR to the promoter region in *E. coli* MG1655 (26). Further details on the deletion and transversion strains are given in Ref. (26). The AcrB^−^ strain was derived from the Keio collection JW0451 (BW25113 *ΔacrB::kan*) and we removed the kanamycin resistance marker following the protocol in Ref. (27). For strains where we inserted the chromosomal fluorescent protein marker, we replaced the *galK* gene with *sfgfp* driven by a strong constitutive promoter.

### Minimum inhibitory concentration

Overnight cultures of WT, MarA^−^, MarA^+^, and AcrB^−^ were diluted 1:100 in LB medium. Diluted cultures were incubated at 37°C with 200 rpm shaking for 4 hours. Cells were then incubated with carbenicillin (0 to 50 μg/ml in 2-fold dilutions) at 37°C for 24 hours with 200 rpm shaking. The minimum inhibitory concentration was determined as the concentration of antibiotic where no visible growth was observed.

### Bacterial growth conditions

For all growth experiments, overnight cultures inoculated from a single colony were diluted 1:500 in LB medium. The diluted cultures were then pre-cultured for 3 hours at 37°C with shaking at 200 rpm. We refer to the end of this pre-culture as time point t = 0 hours. Salicylate was added at t = 0 hours when required, we then continued culturing at 37°C with 200 rpm shaking. OD_660nm_ readings were taken using a BioTek Synergy H1m plate reader each hour.

### Competition experiments

For the competition experiments, each of two strains were cultured overnight separately, then diluted 1:1000 in fresh LB medium. Competition co-cultures were created by mixing cultures of each of two strains together in equal proportion. Co-cultures were then precultured for 3 hours, then at t = 0 salicylate was added where required, as in Ref. (1). OD_660nm_ readings were taken each hour using the plate reader. In addition, immediately following OD_660nm_ measurements, 5 μl of the sample was diluted into 200 μl sterile phosphate-buffered saline (PBS) and we measured fluorescent protein expression with a Guava easyCyte HT flow cytometer. Flow cytometry data were analyzed with custom Matlab scripts. Control experiments using strains without green fluorescent protein expression were carried out to determine the threshold for strain classification (Fig. S1).

### Antibiotic killing experiments

Cultures for antibiotic killing experiments were created by diluting overnight cultures of each of the strains at the percentages shown in Fig. 4. Overnight cultures of each strains were first diluted 1:1000 in fresh LB medium, pre-cultured for 3 hours at 37°C with 200 rpm shaking, then strains were mixed together at percentages as specified in the figure axis. 50 μg/ml of carbenicillin was then added to the co-cultures at t = 0. OD_660nm_ readings were taken using the plate reader four hours after carbenicillin exposure to determine antibiotic killing. Cells were then centrifuged and resuspended in PBS to wash out the antibiotics. Washed cultures were diluted and plated on LB agar for 24 hours in order to determine the number of colony-forming units after antibiotic exposure.

### Dose-response experiments

Overnight cultures of the WT strain were diluted 1:50 in LB medium with 0, 0.6, 1.2, and 5mM salicylate. Diluted cultures with various concentration of salicylate were then pre-cultured for 3 hours at 37°C with 200 rpm shaking. Cells were then incubated with carbenicillin (0 to 50 μg/ml in 2-fold dilutions) at 37°C for 4 hours with 200 rpm shaking. OD_660nm_ readings were taken using the plate reader four hours after carbenicillin exposure to determine cell density.

### marA and acrAB transcriptional reporters

We used *marA* and *acrAB* transcriptional reporters in the WT strain to measure their transcriptional activity upon salicylate exposure. As described in Ref. (28), the promoter region of each gene was cloned upstream of the gene for cyan fluorescent protein (*cfp*) on a low-copy (SC101 origin) plasmid. Salicylate was added at t = 0. The reporter fluorescence level was measured using microscopy after 3 hours of salicylate exposure. Fluorescence distributions were calculated using kernel density.

### Gompertz model

All growth curves were fitted to the modified Gompertz model (Eq. 1) described in Ref. (29). Carrying capacity (A), growth rate (μ), and lag-time (λ) were determined by least-squares fitting to the mean of the experimental data using a custom Matlab script.

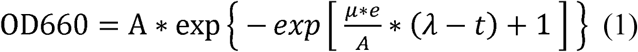

### Selection coefficient

The selection coefficients were calculated using the regression model (Eq. 2) described in Ref. (30) after 4 hours of competition with or without exposure to salicylate.

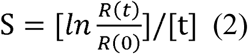

S is the selection coefficient; R(t) is the final competition ratio of mutants to the WT strain, R(0) is the initial competition ratio of mutants to WT strain; t is number of generations in the competition experiment, corresponding to the number of doublings of OD_660nm_ observed (31).

### Lotka-Volterra model

The modified Lotka-Volterra equations (Eqs. 3 & 4) from Ref. (32) were used to simulate competitive growth in the co-culture:

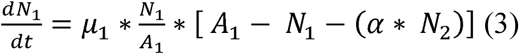

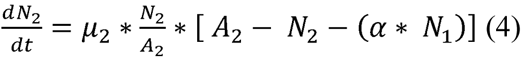

N_i_ is the cell density of strain i, μ_i_ is the maximum growth rate, A_i_ is the carrying capacity, α is the interaction matrix. We assumed no interaction between strains, α = 1. Both μ_i_ and A_i_ were determined using single strain experimental growth curve data fit to the Gompertz model, as described above. Long-time simulations for Fig. 5 assume dilution every 8 generations to the initial cell density, modeling serial transfers. We assume that the generation time is 30 minutes in the laboratory environment. All simulations were conducted using custom Matlab scripts.

## Results and Discussion

### MarA-mediated antibiotic resistance is costly

In order to investigate MarA-mediated resistance, we employed three strains of *E. coli*: one where MarA and its homologs Rob and SoxS are knocked out (denoted MarA^−^) (33), wild type (WT), and a strain where MarA is over-expressed via mutations in the MarR binding sites on its own promoter (MarA^+^). To verify that these strains exhibit differences in antibiotic resistance, we first measured the minimum inhibitory concentration (MIC) of carbenicillin. As expected, we found that the MIC increases as MarA expression levels increase (Fig. S2).

To quantify the cost of MarA-mediated resistance, we measured the growth of MarA^−^, WT, and MarA^+^ strains in LB medium without antibiotics and used fits to the Gompertz model (Eq. 1) to extract parameters associated with growth from the data: growth rate (µ), lag time (*λ*), and carrying capacity (A) (29) (Fig. 1a). We found that of the three model parameters only the growth rate was strongly dependent on MarA expression levels (Fig. 1b), and effects on lag time and carrying capacity were minimal (Fig. S3).

**Figure 1.**
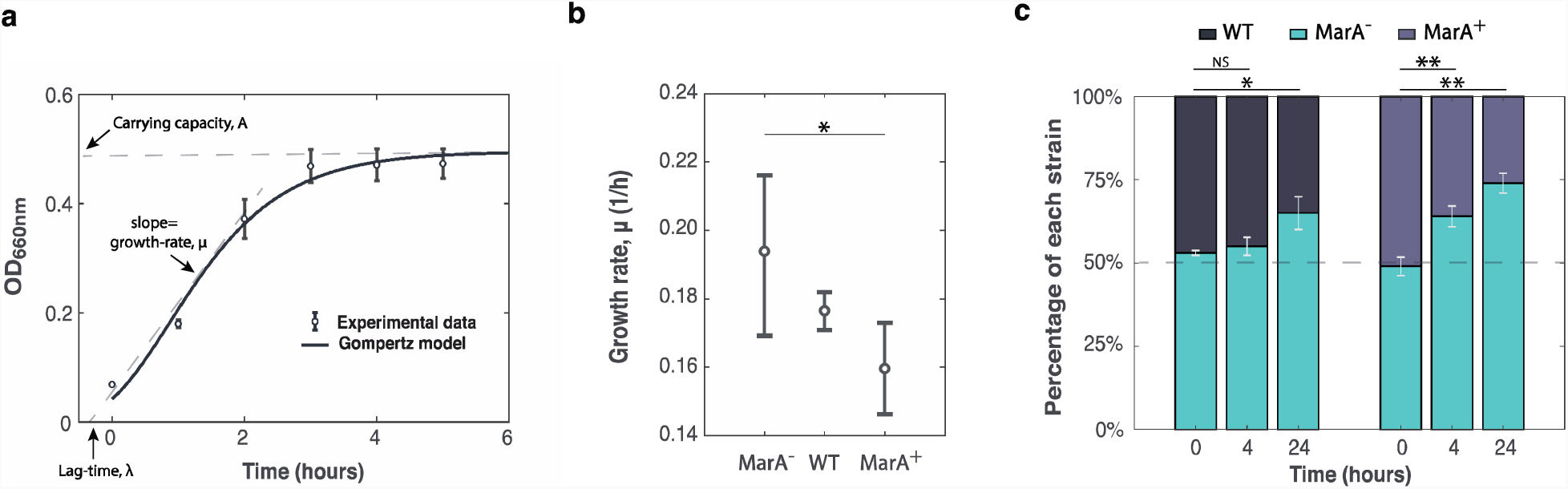
MarA-mediated resistance is costly. (a) MarA^−^ growth curve with experimental and modeling data. Black solid line is a fit to the Gompertz model. Data points show experimental data from six biological replicates. Error bars show standard deviation. Three parameters are extracted from the Gompertz model: growth rate (*µ*), lag time (*λ*), and carrying capacity (A). **(b)** Growth rate extracted from the Gompertz model fits for each strain. Data points show mean values and standard deviation from at least four biological replicates. Lag time and carrying capacity values are shown in Fig. S3. **(c)** Fraction of cells of each strain over time after competition between MarA^−^ and either WT or MarA^+^, with initially equal proportions of the competing strains in well-mixed liquid cultures. Relative proportions were obtained using counts of fluorescent cells from flow cytometry. Error bars show standard deviation from three biological replicates. NS (Not Significant); *P* >0.05; \**P* <0.05; \*\**P* <0.01; \*\*\**P* <0.001, Student’s t-test.

To evaluate whether these differences in growth rates could alter the proportion of bacteria with different MarA expression levels in a mixed population, we carried out competition assays between MarA^−^ and either WT or MarA^+^ strains under antibiotic-free conditions (Fig. 1c). To visualize the differences in the strains we integrated green fluorescent protein (*sfgfp*) into the genome of MarA^−^, and quantified the compositions of the mixed populations by flow cytometry (Methods, Fig. S1). With initially equal representation between the strains, we found that the MarA^−^ strain outgrew MarA^+^ but not WT in a short time window (4 h). In a longer competition (24 h), the MarA^−^ strain outcompeted both WT and MarA^+^ as expected due to the lack of burdensome MarA expression. The change in population composition is strongest between the antibiotic susceptible (MarA^−^) and antibiotic resistant (MarA^+^) strains (Fig. 1c), suggesting fitness differentials are amplified if MarA is overexpressed. As a control, we also competed WT cells with and without genomically integrated *sfgfp* and found that the strains showed equal fitness, as expected (Fig. S4). The inverse relationship between MarA-mediated resistance and fitness supports the concept that differences in fitness cost could result in exclusion of resistant bacteria.

### Salicylate increases fitness cost in resistant strains compared to susceptible strains

Since MarA expression is costly, we reasoned that inducing its expression with salicylate should accelerate the process of biasing the population toward strains that lack MarA expression. To test this, we first measured expression from the *marA* promoter induced with various concentrations of salicylate (Fig. S5a). Using transcriptional reporters for *marA* in WT cells, we observed a clear trend of increased expression as the salicylate concentration increased. We then measured the cost of MarA expression induced by salicylate. As expected, increasing salicylate dosages have a clear negative impact on the growth of WT cells (Fig. 2a). However, the salicylate growth reduction effect can also be observed in the MarA^−^ strain (Fig. 2b), which has *marA* and its homologs deleted. This suggests that salicylate can also reduce growth in a MarA-independent manner, possibly due to global effects such as by inhibiting expression of ATP synthase (34), decreasing membrane potential (35), or reducing metabolic activity (26).

**Figure 2.**
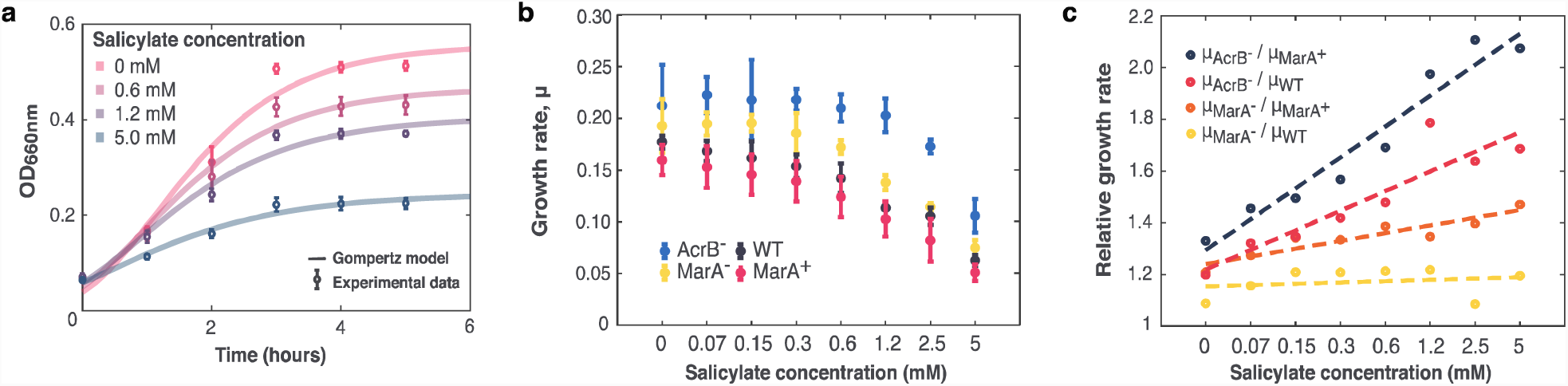
Salicylate increases fitness cost of resistant strains. (a) Growth curves of WT strain under increasing salicylate concentrations. Solid lines show Gompertz model fits. For experimental data, error bars show standard deviation from four biological replicates. **(b)** Growth rates extracted from the Gompertz model for different strains at a range of salicylate concentrations show strain-specific trends between growth rate and salicylate concentration. Data points and error bars show experimental data from at least four biological replicates. **(c)** Relative growth rates between strains at increasing salicylate concentrations. Data points show ratio of mean growth rate between two strains. Dashed lines show linear least squares fit. Slope and its 95% confidence interval (in parenthesis) of the linear fit are: 0.1196 (0.0958, 0.1433) for μ_AcrB_^−^/μ_MarA_^+^;0.0760 (0.0500, 0.1021) for μ_AcrB_^−^/μ_WT_; 0.0300 (0.0201, 0.0398) for μ_MarA_^−^/μ_MarA_^+^; 0.0051 (−0.0125, 0.0226) for μ_MarA_/μ_WT_.

Since salicylate can increase antibiotic resistance and decrease cell growth, in order to measure the costs and benefits of salicylate induction, we further measured the dose-response curve between carbenicillin and salicylate in the WT strain (Fig. S6). We observed a severe reduction in growth at high concentrations of salicylate in the absence of antibiotic, indicative of costs imposed by salicylate exposure. However, the cells have a growth benefit from salicylate induction when the carbenicillin concentration is high, suggesting a trade-off between benefits and costs of salicylate induction.

We next asked whether salicylate could increase the cost in strains with MarA expression (WT and MarA^+^) relative to the MarA^−^ strain. To do this, we compared the relative growth rates between MarA^−^ with either WT or MarA^+^, measured by the ratio of the growth rates between the two strains in increasing concentration of salicylate (Fig. 2c). Surprisingly, we found a weak positive correlation between relative growth rate and salicylate concentration in MarA^−^ vs. MarA^+^, but not in MarA^−^ vs. WT. Note that MarA^+^ cannot respond to salicylate via the native repressor MarR due to mutations in its MarR binding sites, indicating that this response to salicylate involves factors beyond MarA overexpression, potentially through induction of the MarA homolog Rob (33). We also observed similar effects using fitting to a Hill function (1) to quantify the growth cost in different strains as a function of salicylate concentration (Fig. S7).

### AcrAB-TolC efflux pump is a major contributor to MarA-mediated resistance burden

We next asked where the cost of salicylate-induced MarA burden was coming from. MarA regulates many downstream genes, but the AcrAB-TolC efflux pump is known to play a critical role in resistance and imposes a significant energy cost (3). A transcriptional reporter for *acrAB* also showed that *acrAB* transcription can be induced by salicylate (Fig. S5b). To test its role in salicylate-induced burden, we constructed AcrB^−^, an *acrB* knockout strain that renders the efflux pump nonfunctional. We also observed lower MIC values in AcrB^−^ compared to WT or MarA^+^ (Fig. S2).

We measured growth rate of AcrB^−^ with MarA^+^ and WT and compared it under various concentration of salicylate (Fig. 2b). Interestingly, salicylate has only a modest effect on the growth rate of AcrB^−^ below 2.5 mM concentrations. We also observed a positive linear relationship between salicylate and the relative growth rate of AcrB^−^ with WT and MarA^+^ (Fig. 2c). These results demonstrate that salicylate imposes less burden if the efflux pump is deleted. It is important to note that the AcrB^−^ strain has intact copies of MarA and its homologs. Thus, AcrB^−^ can still respond to salicylate exposure through MarA and its homologs, but the cost will not be increased in low salicylate concentrations if AcrAB-TolC cannot be produced. These results suggest that salicylate can induce much higher burden in the resistant strain (MarA^+^) than in the susceptible strain (MarA^−^), and this burden is mainly contributed by the AcrAB-TolC efflux pump.

### Salicylate accelerates competitive exclusion of resistant bacteria

We next sought to test whether using salicylate to increase the relative cost in the resistant strain (MarA^+^) accelerates its exclusion in competitive environments. Since we found AcrAB-TolC to be largely responsible for the salicylate induced fitness cost, we first measured competitive exclusion between AcrB^−^ and MarA^+^. Using an AcrB^−^ strain with genomically integrated *sfgfp*, we measured the fraction of AcrB^−^ and MarA^+^ cells by flow cytometry and the growth of the competition culture using a plate reader.

In the absence of salicylate, small differences in growth rates are enough to allow the susceptible strain AcrB^−^ to out compete the resistant strain MarA^+^, although the exclusion effect is not large (Fig. 3a). When the salicylate dosage was raised to 0.6 mM, the fraction of AcrB^−^ cells increased because salicylate reduces the growth of the resistant strain MarA^+^ without imposing a significant impact on the susceptible strain AcrB^−^ (Fig. 3b). This difference is further magnified at higher levels of salicylate exposure (Fig. 3c).

**Figure 3.**
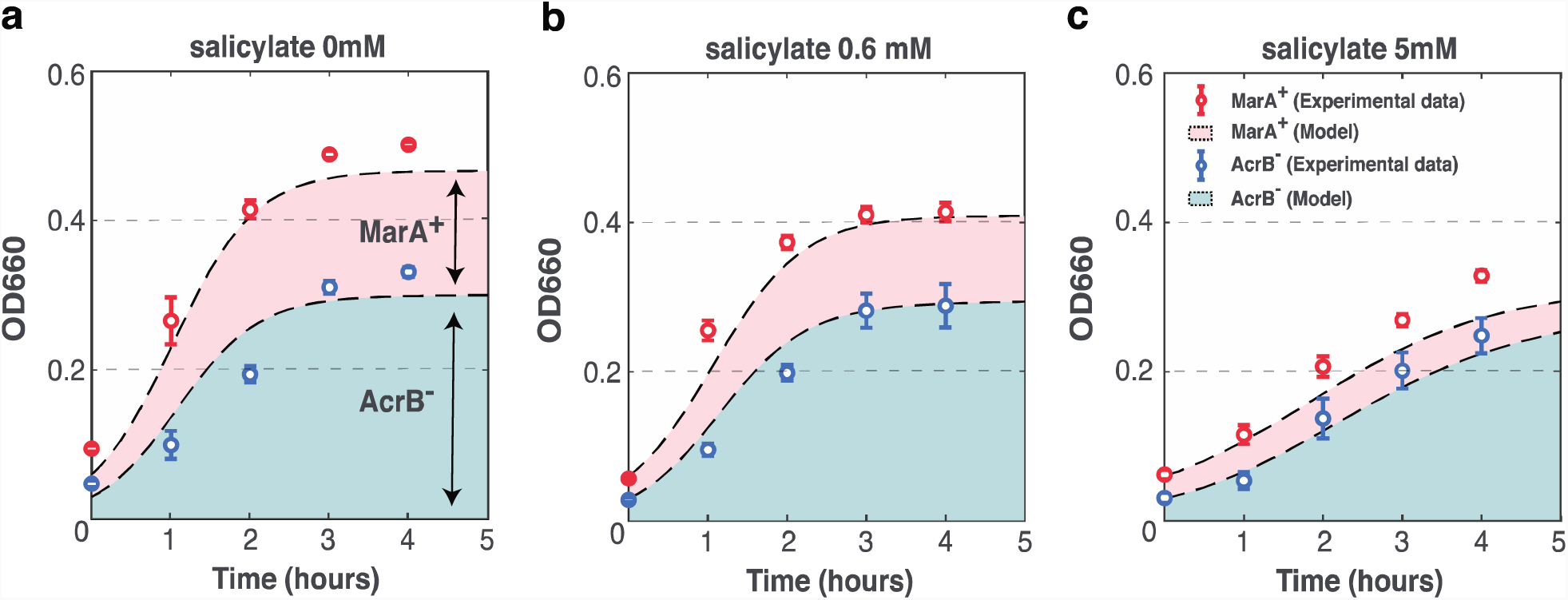
Salicylate accelerates competitive exclusion of resistant bacteria. Competitions of AcrB^−^ (blue) with MarA^+^ (red) in **(a)** 0 mM **(b)** 0.6 mM, and **(c)** 5 mM salicylate seeded with initially equal populations. The red and blue shaded areas represent the predicted growth of each subpopulation using the Lotka-Volterra model, with the total area representing the combined population growth. Data points show experimental mean values and standard deviations from three biological replicates.

To predict the competition dynamics between strains in different concentration of salicylate, we employed a mathematical model that we fit to the single-species growth rate (μ) and carrying capacity (A) data and used it to predict two-species competition under various salicylate conditions. The growth rates and carrying capacities calculated from single culture growth curves fit using the Gompertz model were applied to a Lotka-Volterra model for competitive growth (Eqs. 2-3). The model shows good agreement with the experimental data in the dynamics of both growth and the population fraction for each strain (Fig. 3).

We observed similar effects with salicylate when competing WT with AcrB^−^ and MarA^−^ (Fig. S8). However, competition between WT and MarA^+^ did not lead to competitive exclusion. Since salicylate can significantly decrease bacterial growth (Fig. 2b), we verified that our results were not an artifact of differences in growth rate by calculating the selection coefficients during competition between WT and all mutants under salicylate exposure (Fig. S9). The selection coefficient takes into account growth differences at higher salicylate concentrations (30). We observed similar trends in selection coefficients and results with cell counts, where AcrB^−^ and MarA^−^ have significantly higher finesses compared with WT at high concentrations of salicylate.

### Reducing population fraction of resistant cells increases susceptibility to antibiotic exposure

To study the impact of the fraction of resistant strains on antibiotic resistance of a population, we subjected mixed populations with different proportions of susceptible (AcrB^−^) and resistant (MarA^+^) strains to a lethal dose of carbenicillin (50 *µ*g/mL) for four hours and then measured the optical density and colony-forming units (CFU). The results show that carbenicillin resistance is directly related to the fraction of MarA^+^ cells in the culture (Fig. 4). We found similar results in mixed populations with MarA^+^ and MarA^−^ (Fig. S10) and MarA^+^ and WT (Fig. S11c), but not in populations with AcrB^−^ and WT or MarA^−^ and WT (Fig. S11a-b). Decreasing the proportion of MarA^+^ can significantly reduce the antibiotic resistance of the population. This result supports our hypothesis that antibiotic resistance levels decrease when a susceptible strain out-competes a resistant strain.

**Figure 4.**
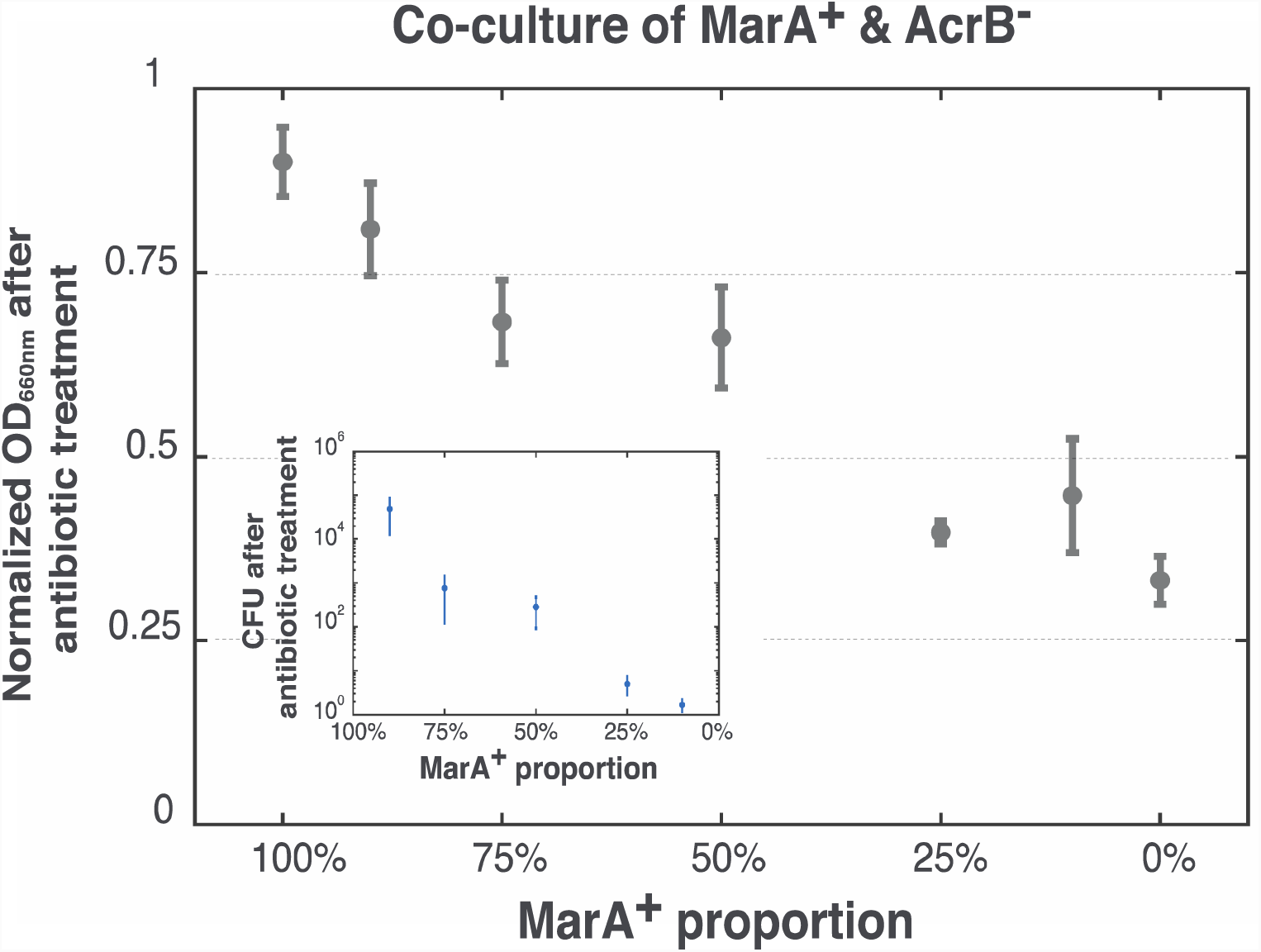
Reducing fraction of resistant cells increases susceptibility to antibiotic killing. Optical density of co-cultures of sensitive (AcrB^−^) and resistant (MarA^+^) strains after applying 50 *µ* g/mL carbenicillin for 4 hours compared to optical density before treatment. Data points show mean values and standard deviation from three biological replicates. Inset figure shows colony-forming units (CFU) of co-cultures of AcrB^−^ and MarA^+^ strains. Data points show mean values and standard deviation from six biological replicates.

### Modelling population shifts under salicylate treatment

In the real-world environment, specific mutants may only constitute a small proportion of cells. To study the population dynamics starting from a very small initial proportion of the susceptible strain, we next used the Lotka-Volterra model to make predictions about the effects of salicylate exposure on mixed populations of AcrB^−^ and MarA^+^ (Fig. 5).

**Figure 5.**
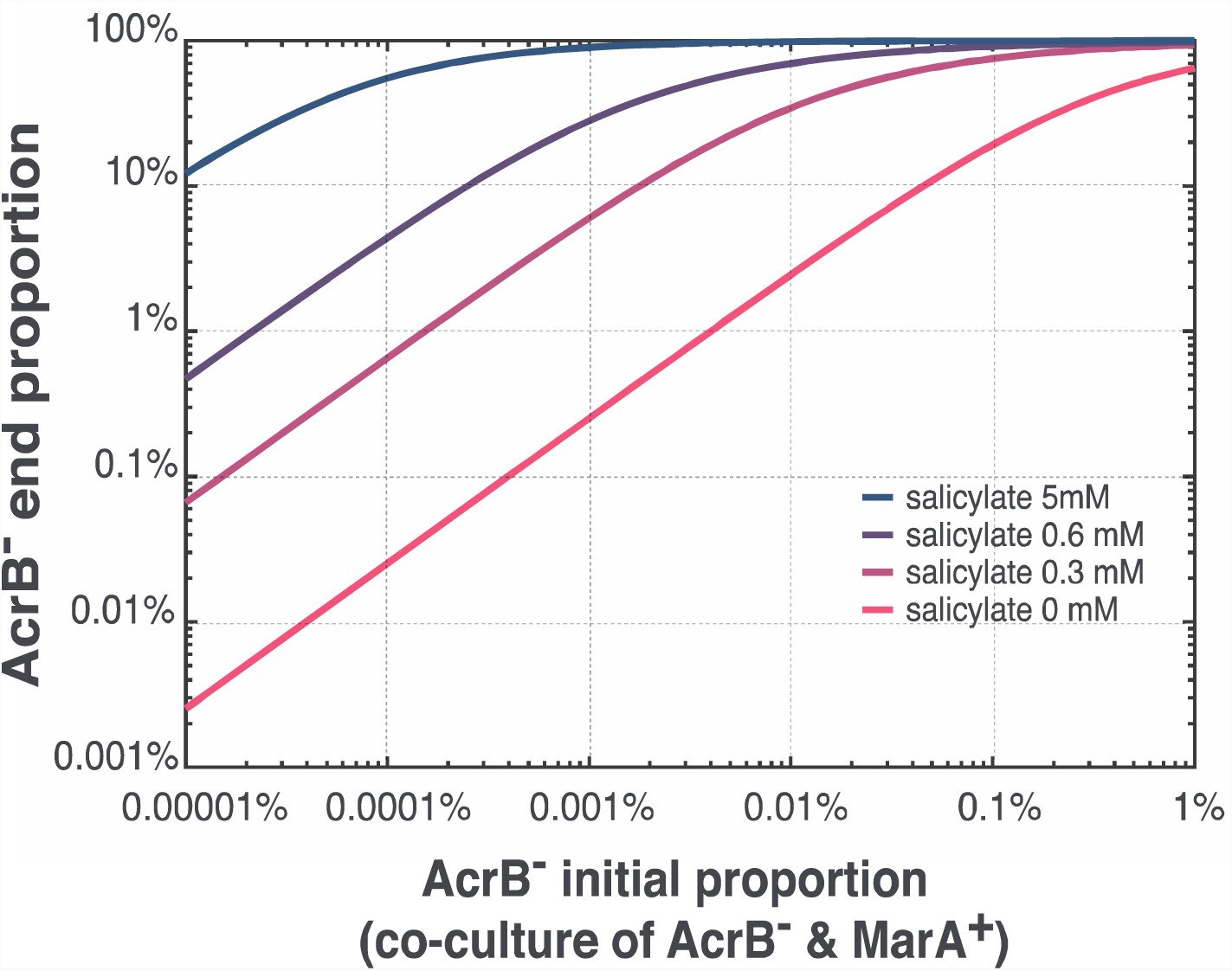
Mathematical modeling predicts salicylate can accelerate population shift towards antibiotic susceptible strains. Predicted population compositions of competitions between AcrB^−^ and MarA^+^ obtained by extending the Lotka-Volterra model over 60 generations. Lines represent the final population fraction of susceptible cells for a particular salicylate concentration.

We simulated competitive exclusion by starting with various initial proportions of the susceptible strain AcrB^−^ and applying the model over the course of 60 generations (approximately 1 week in real time (7)). The model predicts that resistant bacteria (MarA^+^) will be out competed by AcrB^−^ much more rapidly with salicylate exposure than under conditions without salicylate. For example, if the susceptible strain AcrB^−^ initially comprises 0.01% of the population, after 60 generations more than 30%, 80%, and 99% of MarA^+^ is excluded in the presence of 0.3 mM, 0.6 mM, and 5 mM salicylate, respectively. These results are in stark contrast to the conditions without salicylate, where less than 5% of the MarA^+^ can be out-competed by AcrB^−^ with the same initial composition. The model predicts that AcrB^−^ can outgrow MarA^+^ and that salicylate accelerates this process, shifting the population towards antibiotic susceptible strains.

## Conclusions

Antibiotic resistance mechanisms frequently impose a significant fitness cost, suggesting the potential for competitive exclusion of resistant strains. However, this process is usually slow due to downregulation of the resistance machinery in the absence of inducers. One study combining mathematical modeling and experimental data predicted that it would take approximately 1.5 years to reverse tetracycline resistance by replacing 99.9% of the cells carrying the *tetA* resistance gene (7). However, in the natural environment, bacteria commonly interact with other chemicals that could potentially induce expression of resistance genes. Costs imposed by induction could bias population composition.

In this study, we sought to unravel the impact of salicylate exposure on MarA-mediated resistance cost. We establish a mathematical model to predict the population dynamics under different levels of salicylate exposure and extended this to different initial population percentages. We found that salicylate imposes a higher cost in MarA-mediated resistant strains (MarA^+^) than in susceptible (MarA^−^ and AcrB^−^) strains. Competition assays between MarA^+^ and AcrB^−^ showed a positive correlation between salicylate concentration and the relative fitness cost. This suggests that with higher salicylate induction, the relative fitness cost of resistant strains will increase, accelerating their competitive exclusion. Mathematical modelling also predicted mixed population dynamics from single strain growth parameters, and confirmed the effectiveness of salicylate exposure on accelerating the population bias towards susceptible strains. The resulting lower proportion of resistant cells in the population leads to greater lethality to carbenicillin. We further explored the origin of salicylate-mediated cost, and found that the AcrAB-TolC efflux pump contributes to escalated fitness cost in MarA-mediated resistant strains.

Other factors that might affect competitive exclusion include the effective concentrations of salicylate and other environmental chemicals. The physiological concentrations of salicylate in human circulating plasma have been measured and vary between 0.1 μM to 0.5 mM (15) overlapping with values we tested here (Fig. 5). Furthermore, MarA and AcrAB are also involved in resistance to many environmental chemicals beyond antibiotics. Further research may enable understanding of the interplay between salicylate and other chemicals and their combinatorial effects.

Our study indicates the potential for important interaction effects between environmental chemicals and population composition. Understanding the interplay between environmental chemicals and the cost of resistance may help to explain and predict the population dynamics of resistant bacteria in the environment.

## Author Contributions

T.W. and M.J.D. conceived and designed the experiments; T.W. and C.K. performed the experiments and analyzed the data; T.W., C.K., and M.J.D. wrote the manuscript. All authors gave final approval for publication.

## Acknowledgments

We thank members of the Dunlop Lab for helpful discussions. This work was supported by the National Institutes of Health grant 1R01AI102922.

