## Supporting Information for "Salicylate increases fitness cost associated with MarA-mediated antibiotic resistance"

**Supporting Material**

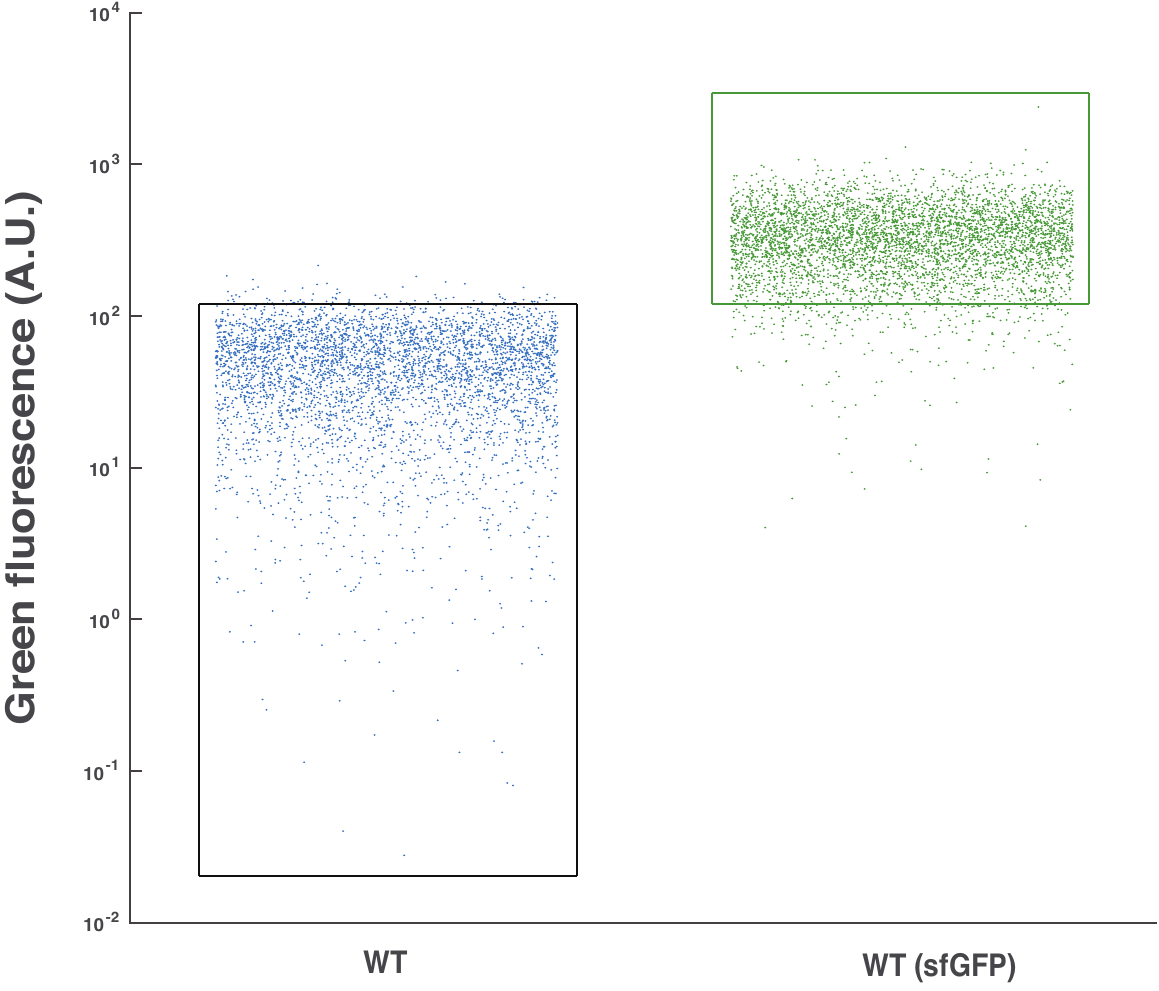

**Figure S1. Green fluorescence level of WT with and without genomically integrated *sfgfp* determined by flow cytometry.** WT strain with and without sfGFP. Each dot represents a green fluorescence reading of one cell in a sample of 5000 cells by flow cytometry. Black and green boxes show thresholds for distinguishing between fluorescent and non-fluorescent strains.

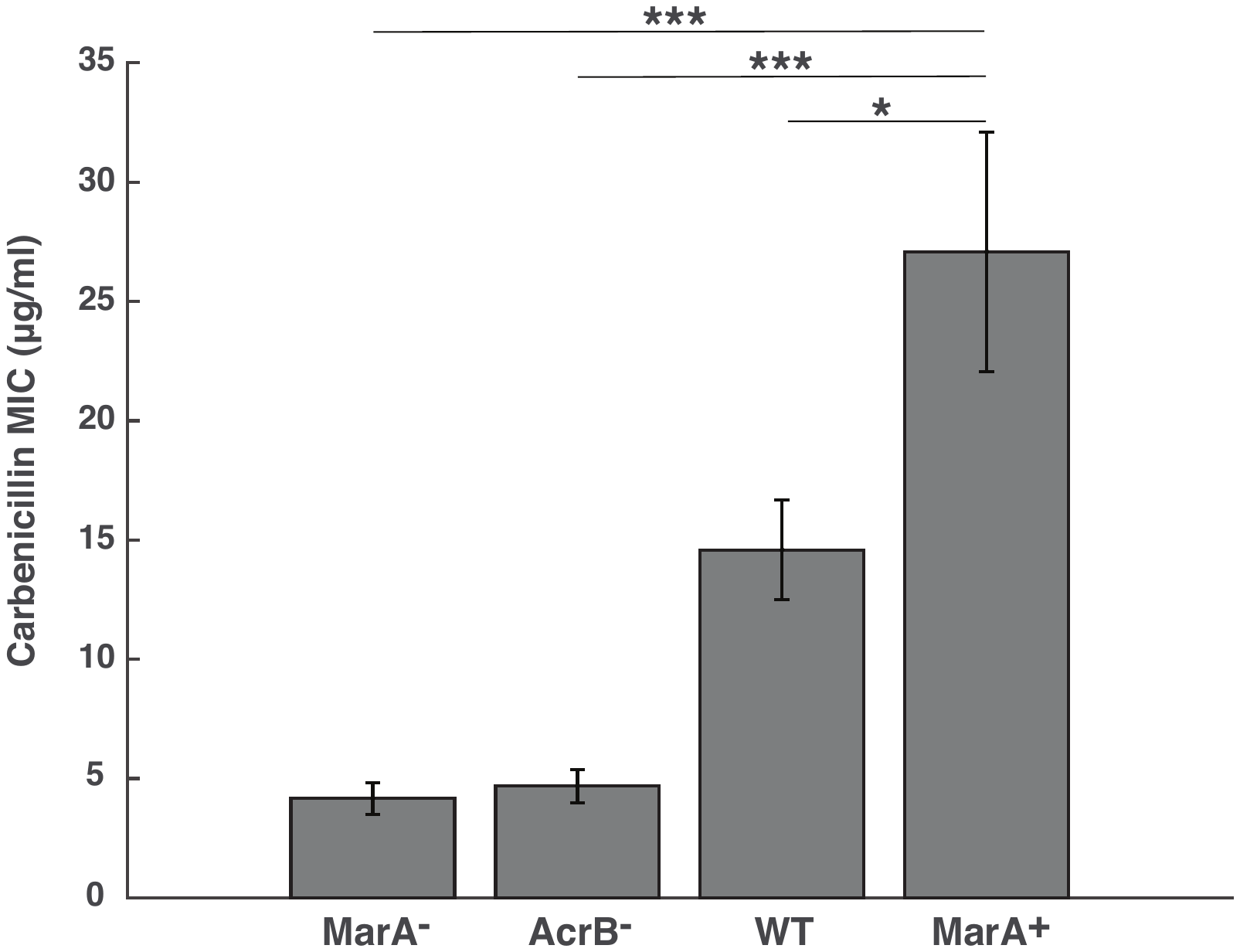

**Figure S2. Minimum inhibitory concentration of carbenicillin in MarA^-^, AcrB^-^, WT, and MarA^+^ strains.** Error bars show standard deviation from six biological replicates. **P*<0.05; ***P*<0.01; ****P*<0.001, Student’s t-test.

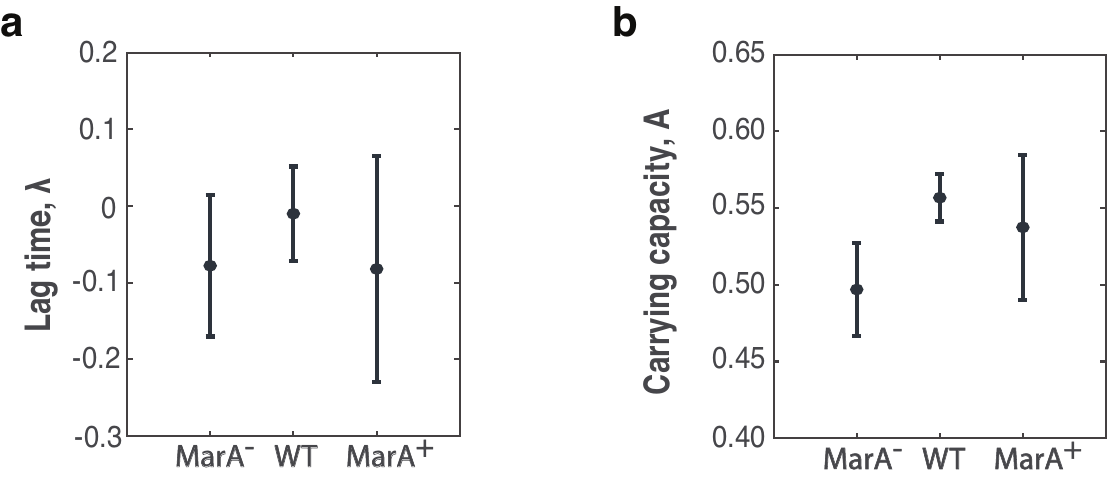

**Figure S3. Lag time (λ) and carrying capacity (A) extracted from Gompertz model fits for each strain.** Data points show mean values and standard deviation from six biological replicates. Differences between the strains are not statistically significant, *P* > 0.05 by a Student’s t test.

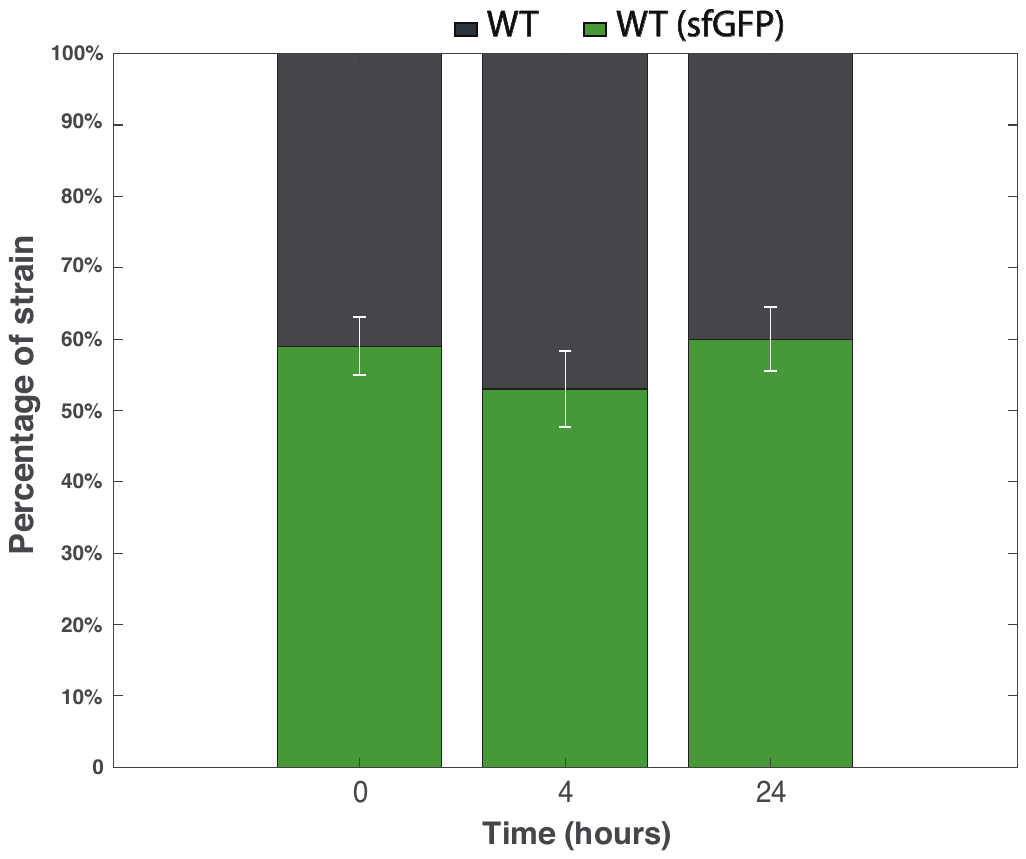

**Figure S4. Competition assay between WT strains with and without genomically integrated *sfgfp.*** Fraction of cells of each strain over time after competition between the WT and WT (sfGFP) with initially equal proportions of the competing strains in a well-mixed liquid culture. Relative proportions were obtained using counts of each fluorescent cell from flow cytometry. Error bars show standard deviation from three biological replicates.

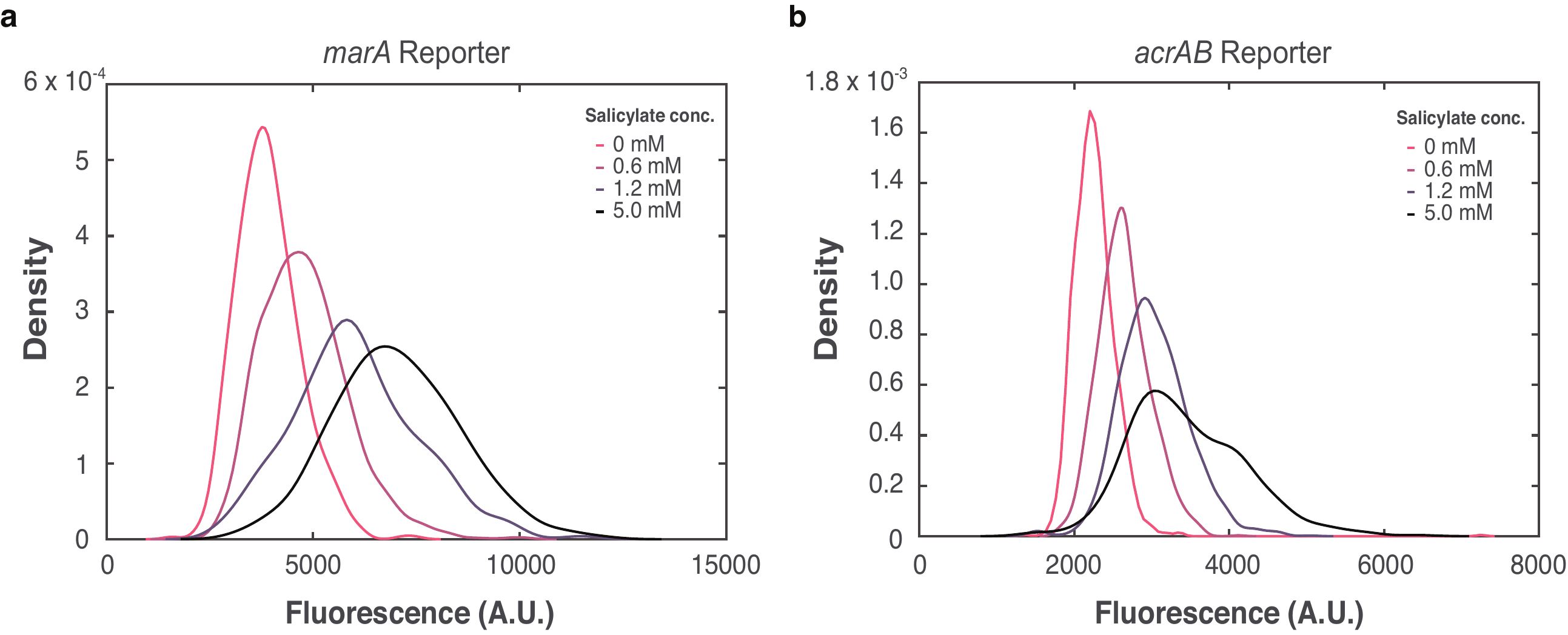

**Figure S5.** **Fluorescence distributions from (a)** *marA* reporter and **(b)** *acrAB* reporter in the WT strain with and without salicylate exposure.

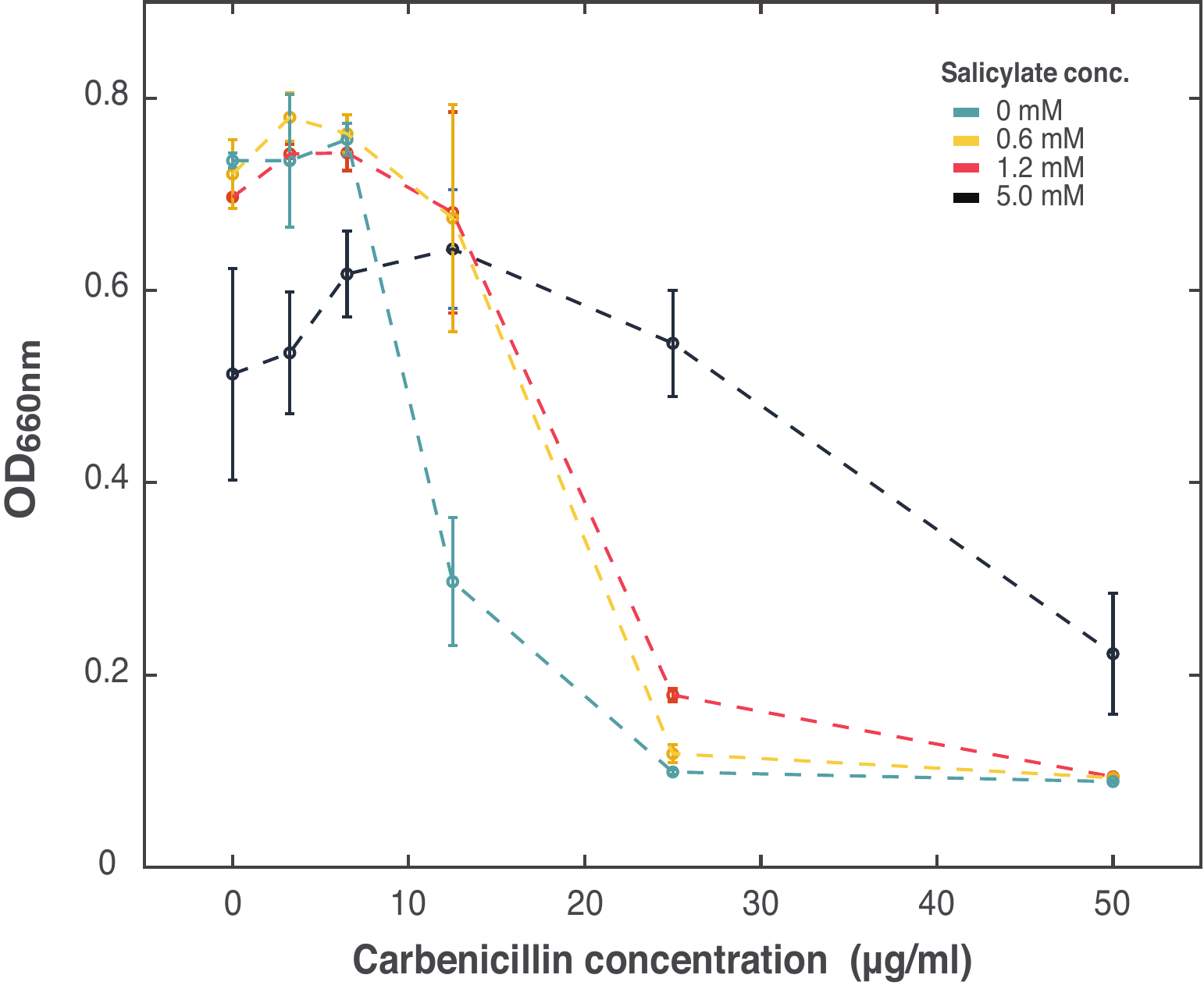

**Figure S6.** **Dose-response curve between carbenicillin and salicylate for the WT strain.** Optical density of WT strain growing in cultures with carbenicillin and salicylate. Data points show mean values and standard deviation from three biological replicates.

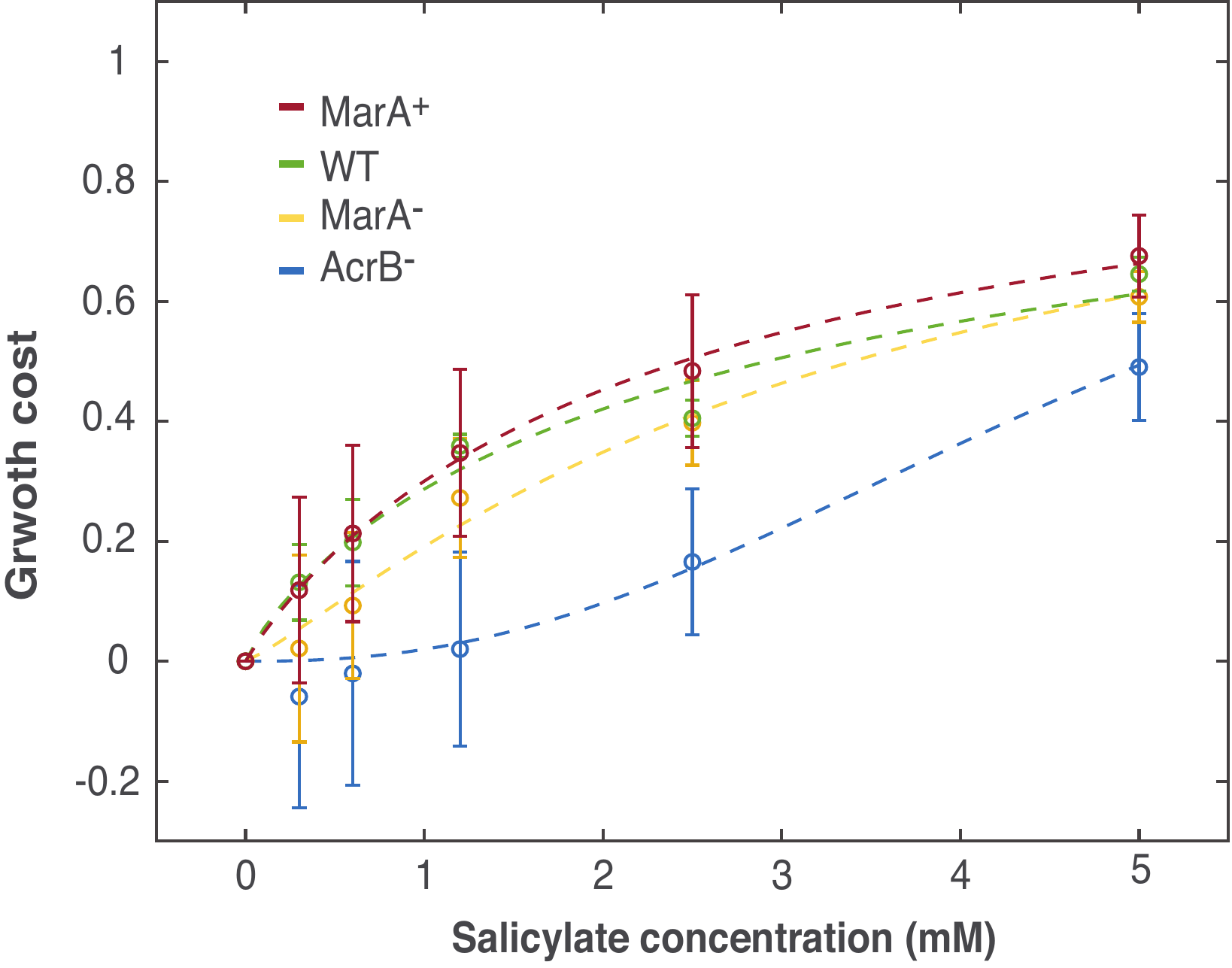

**Figure S7. Growth cost of strains as a function of salicylate concentration.** Growth cost is defined as the reduction of growth rate of cells treated with salicylate relative to untreated cells. Dashed lines are fits to a Hill function h(x) = x^n^/(K^n^+x^n^), where K equals to the salicylate concentration which inhibited cell growth by 50%, as described in Ref (1). K=5.05 (3.45, 5.83) mM, n=2.04 (1.37, 4.26) for the AcrB^-^ strain; K=3.40 (1.49, 4.07) mM, n=1.18 (0.87, 2.71) for the MarA^-^ strain; K=2.92 (1.15, 3.88) mM, n=0.85 (0.55, 1.94) for the WT strain; K=2.44 (1.05, 2.67) mM, n=0.95 (0.85, 2.21) for the MarA^+^ strain. 95% confidence intervals from nonlinear least square fitting are listed in parentheses. Data points and error bars show mean and standard deviation of experimental data from at least four biological replicates.

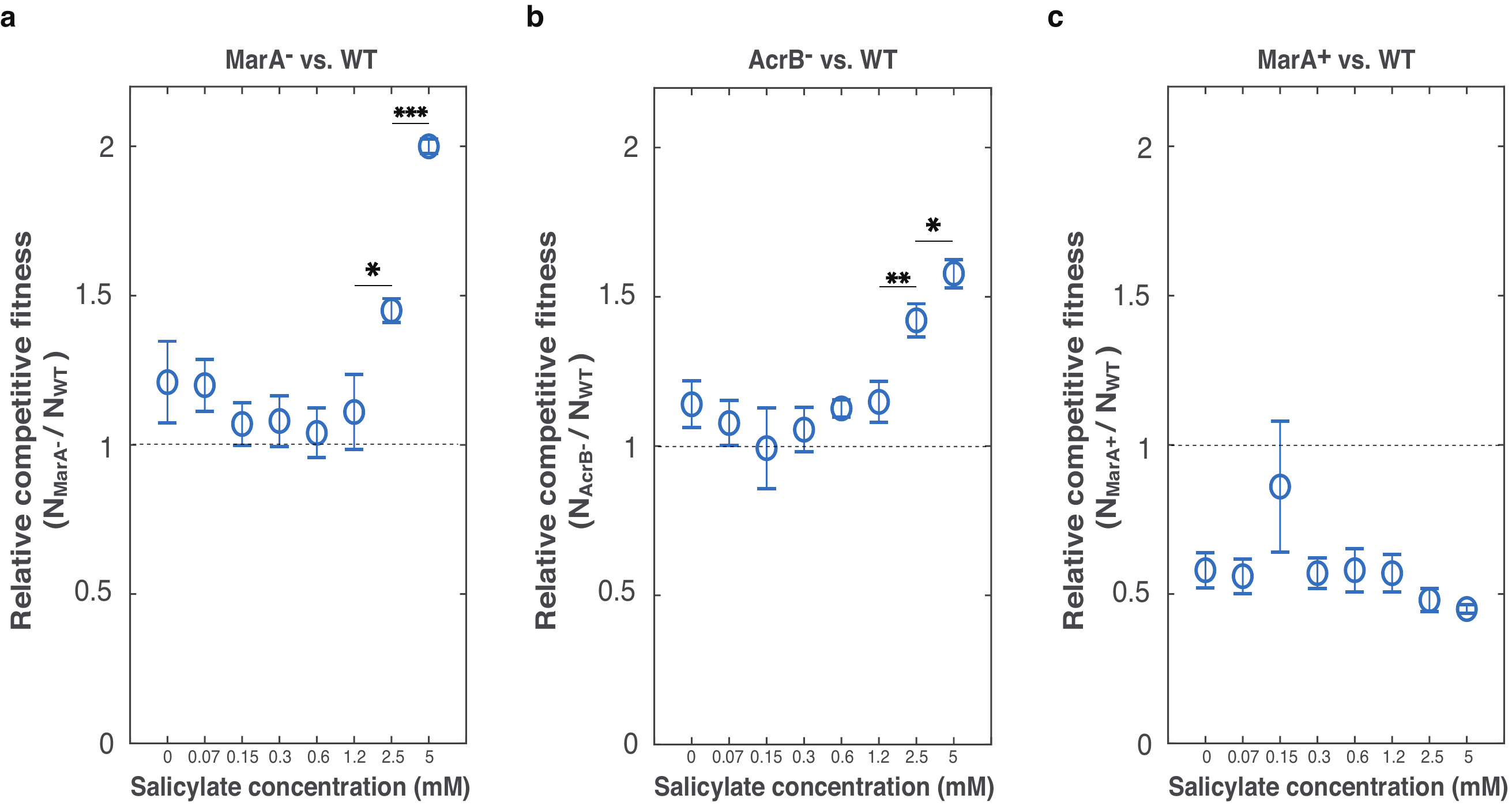

**Figure S8.** **Relative competitive fitness between mutants and WT in various salicylate concentrations.** Competitions of WT with **(a)** MarA^-^**, (b)** AcrB^-^, and **(c)** MarA^+^ in various concentrations of salicylate seeded with initially equal populations. Relative competitive fitness was measured by dividing cell counts of each mutant (N_MarA_^-^, N_AcrB_^-^, or N_MarA_^+^) by cell counts of WT (N_WT_). Cell counts were obtained by counting fluorescent cells from flow cytometry. Strains with genomically integrated *sfgfp* are MarA^-^ in (a), AcrB^-^ in (b) and WT in (c). Data points show mean values and standard deviation from three biological replicates. **P*<0.05; ***P*<0.01; ****P*<0.001, Student’s t-test.

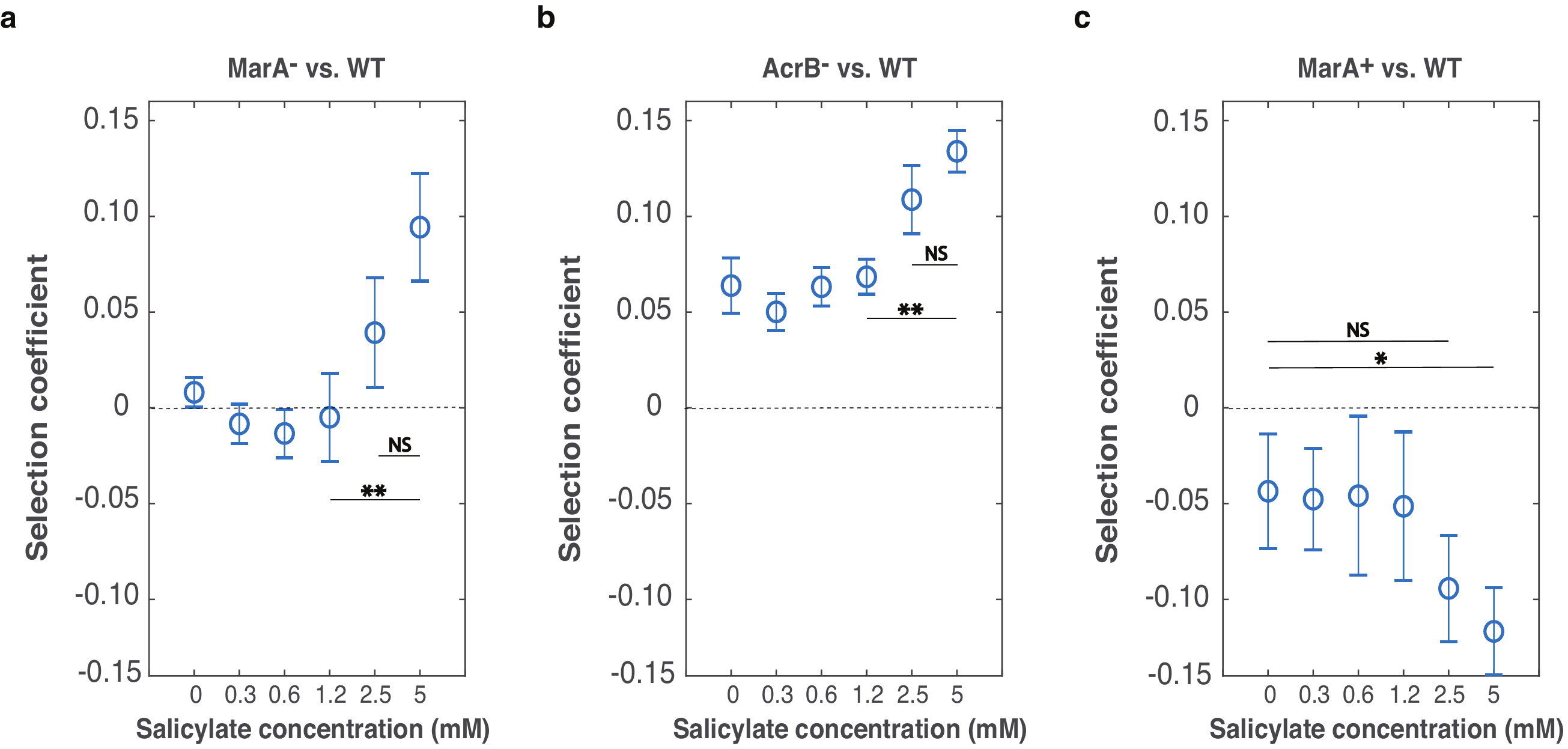

**Figure S9.** **Selection coefficients as a function of salicylate concentration in the competition experiments between WT and (a)** MarA^-^, **(b)** AcrB^-^, and **(c)** MarA^+^. The selection coefficients were calculated using the regression model described in methods section. Strains with genomically integrated *sfgfp* are MarA^-^ in (a), AcrB^-^ in (b) and WT in (c). Data points show mean values and standard deviation from three biological replicates. **P*<0.05; ***P*<0.01; ****P*<0.001, Student’s t-test.

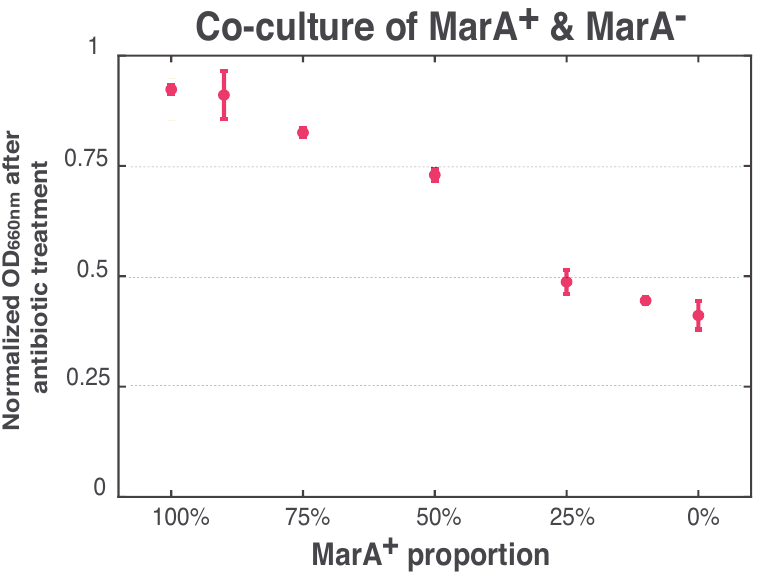

**Figure S10.** **Reducing population fraction of resistant cells increases susceptibility to antibiotic killing.** Optical density of co-cultures of MarA^+^ and MarA^-^ strains after applying 50 $\mu$g/mL carbenicillin for 4 hours compared to optical density before treatment. Data points show mean values and standard deviation from three biological replicates.

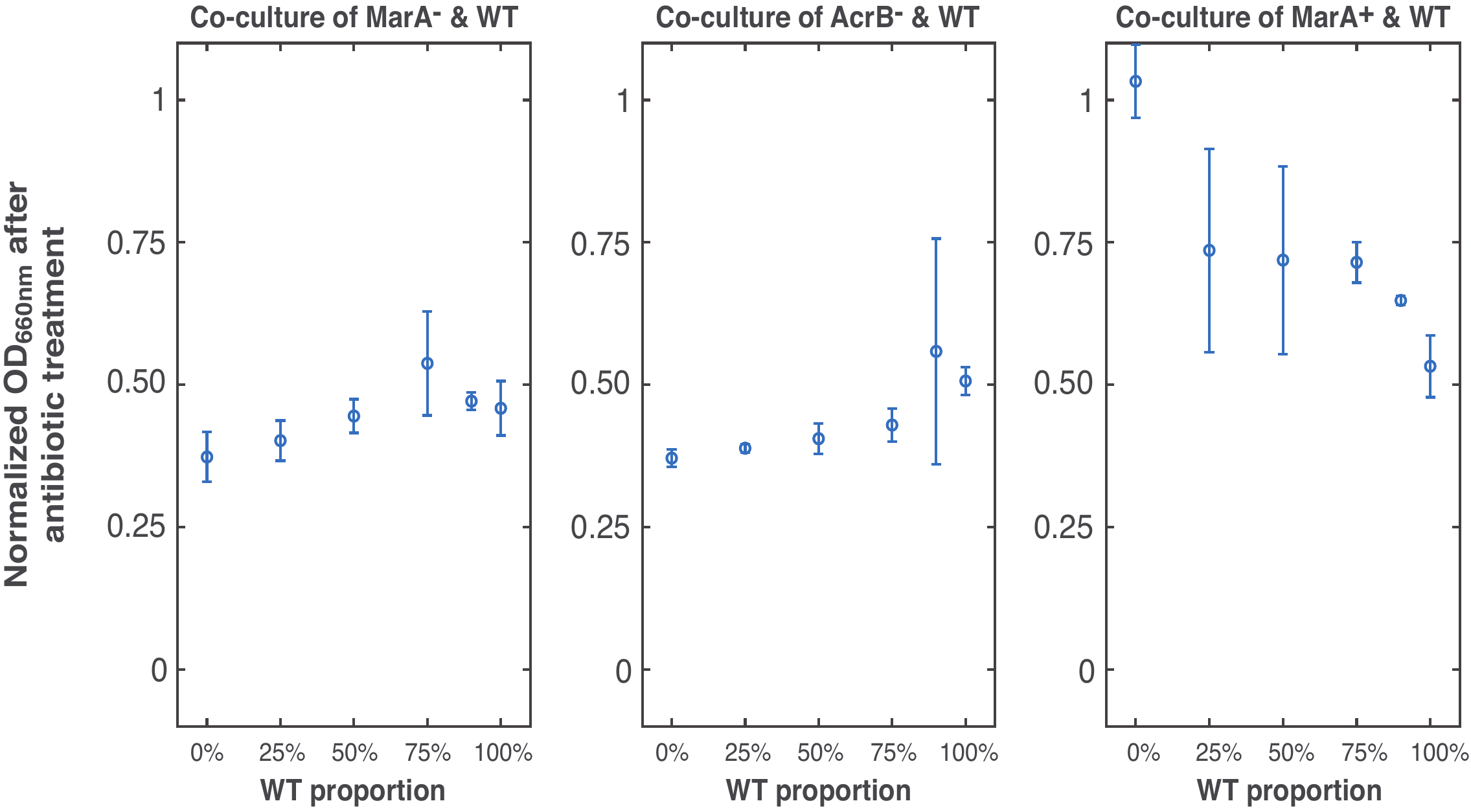

**Figure S11. Normalized OD_660nm_ after 4 hours of carbenicillin treatment in co-cultures of mutants and WT.** Optical density of co-cultures of mutants and WT strains after applying 50 $\mu$g/mL carbenicillin for 4 hours compared to optical density before treatment. Data points show mean values and standard deviation from three biological replicates.
